## Supplementary material for "Oxygen tension regulates lysosomal activation and receptor tyrosine kinase degradation": suppl legend

**Suppl. Fig. 1.** Serum-starved HUVECs and HeLa cells were cultured in either 20% or 1% oxygen in the presence or absence of 50ng/ml of EGF for 5 hrs, followed by Western blot (A). Serum-starved HUVECs were cultured in either 20% or 1% oxygen in the presence or absence of 50ng/ml of VEGF for 5 hrs, followed by Western blot (B). Serum-starved HUVECs were treated with 50 ng/ml of VEGF in the presence or absence of 5 μM of chloroquine (CQ) for 5 hours (C).

**Suppl. Fig. 2.** HeLa cells transfected with Myc tagged TFEB were incubated in 20% $O_2$ in the presence of absence of 10nM of AZD2014, or 1% $O_2$, for 5 hrs, followed by immunostaining for TFEB (A). HeLa cells transfected with Myc tagged TFEB were incubated in 20% or 1% $O_2$ in the presence of absence of 2 μM of MHY1485 for 4 hrs, followed by immunostaining for TFEB (green) and counter stained for wheat germ agglutinin (red) (B). Images were collected from confocal microscopy. Representative images were shown.

**Suppl. Fig. 3.** HeLa cells transfected with TFEB-myc expression vectors for 1 day, followed by stimulation with vehicle, 2 μM of MHY1485 or 10nM of AZD2014 for 30 minutes. TFEB was immunoprecipitated with antibodies against the Myc tag, followed by mass-spectrometry analysis.

**Suppl. Fig. 4.** HeLa cells transfected with WT, S463A or S463D TFEB constructs were incubated in either 20% or 1% oxygen in the presence of absence of 2 μM of MHY1485 for 4 hrs, followed by immunostaining for TFEB. Images were collected from confocal microscopy. Representative images were shown.
