## Supplementary figures and images for "Oxygen tension regulates lysosomal activation and receptor tyrosine kinase degradation"

### suppl fig

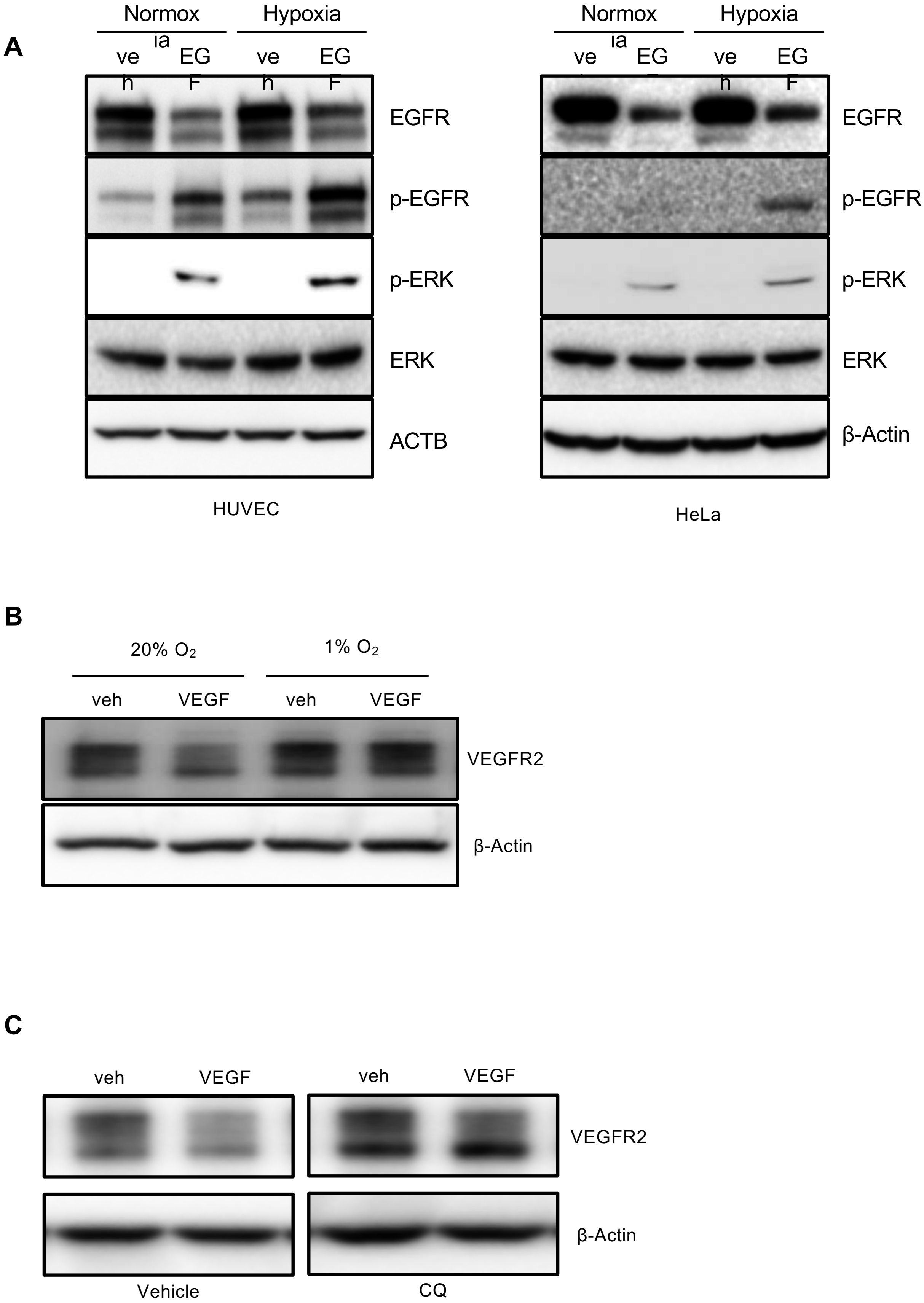

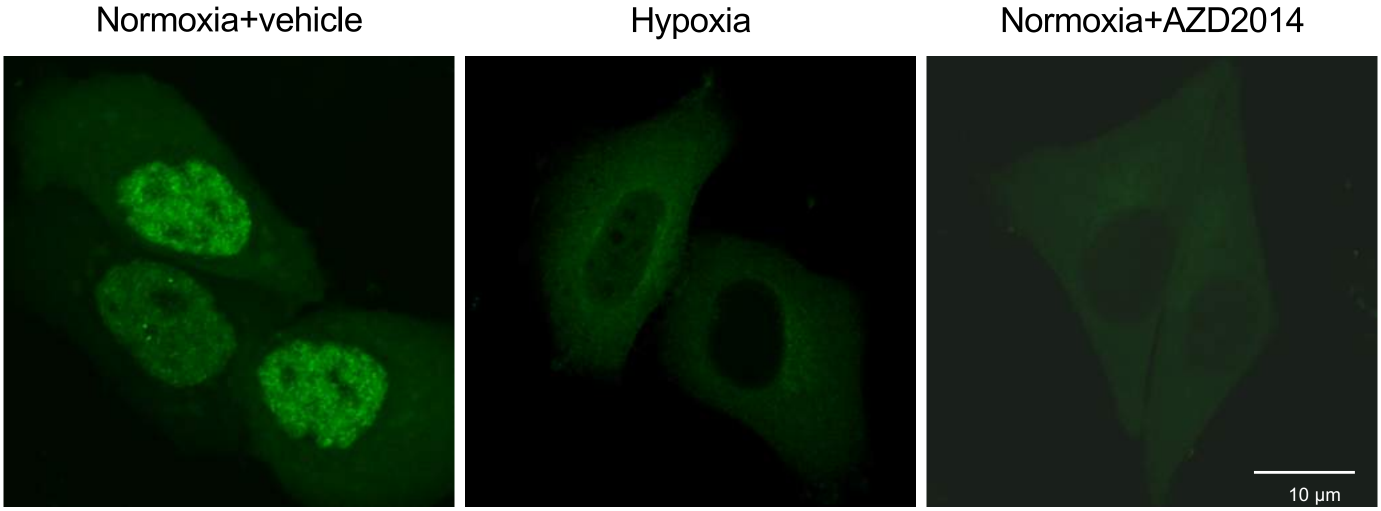

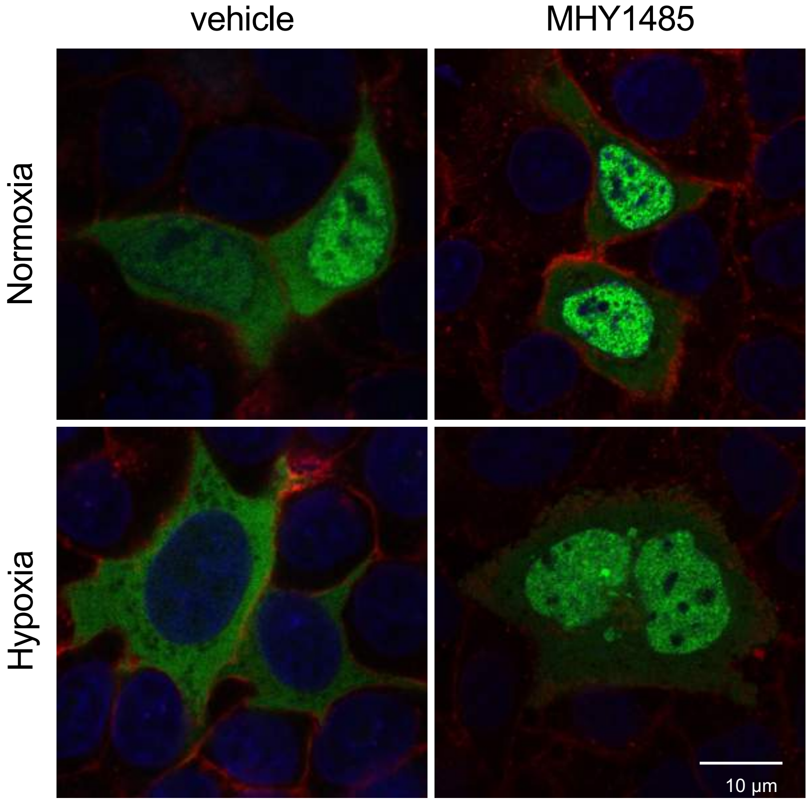

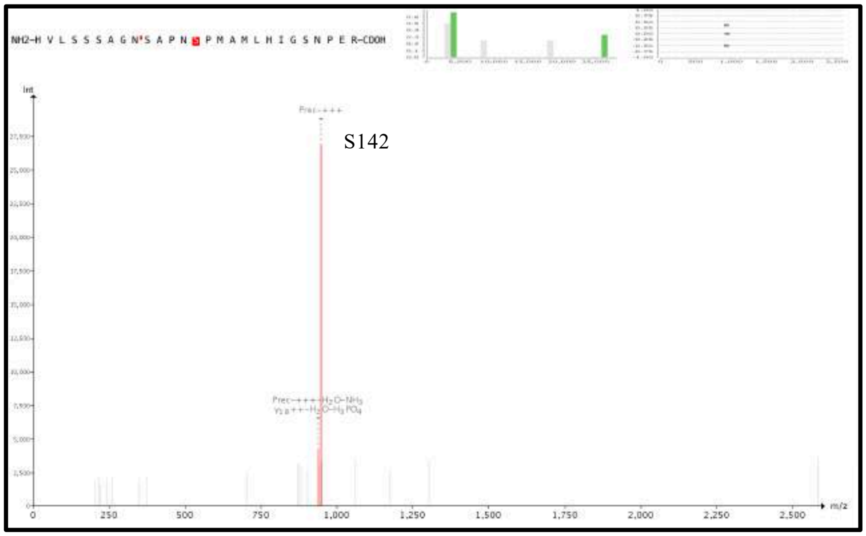


Vehicle

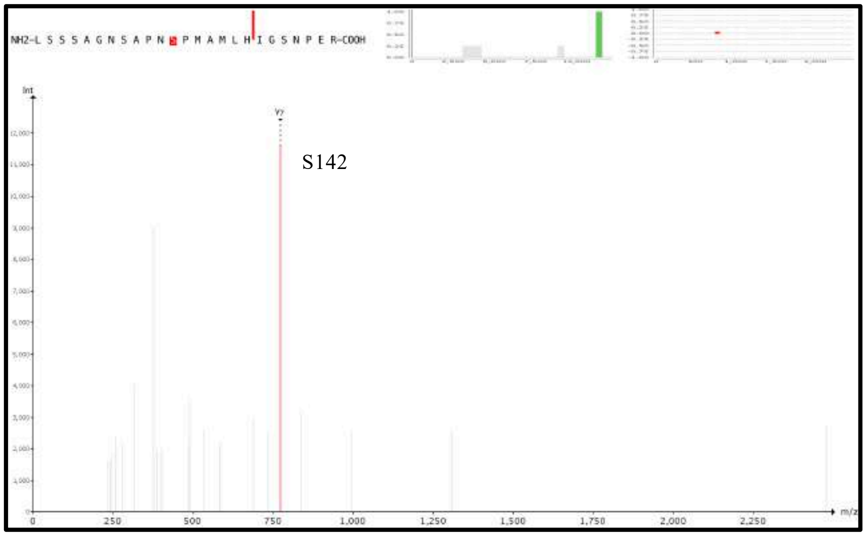


MHY1485

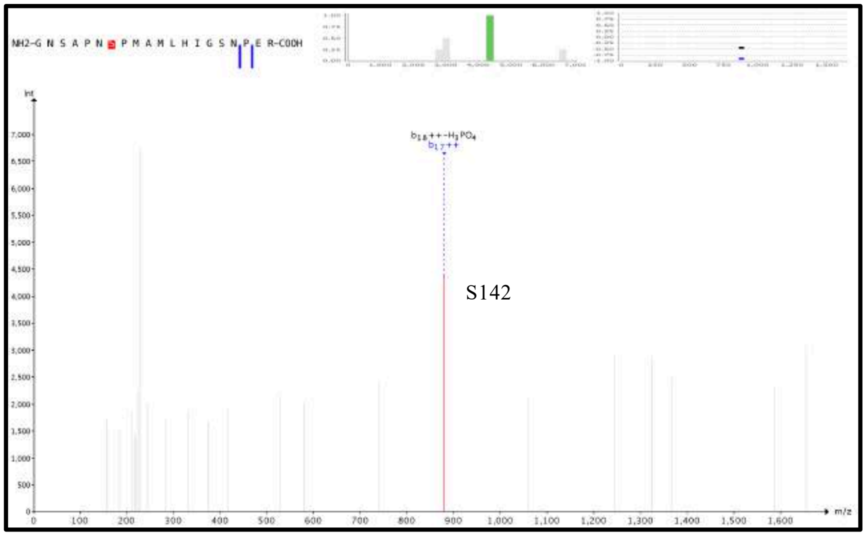


AZD2014

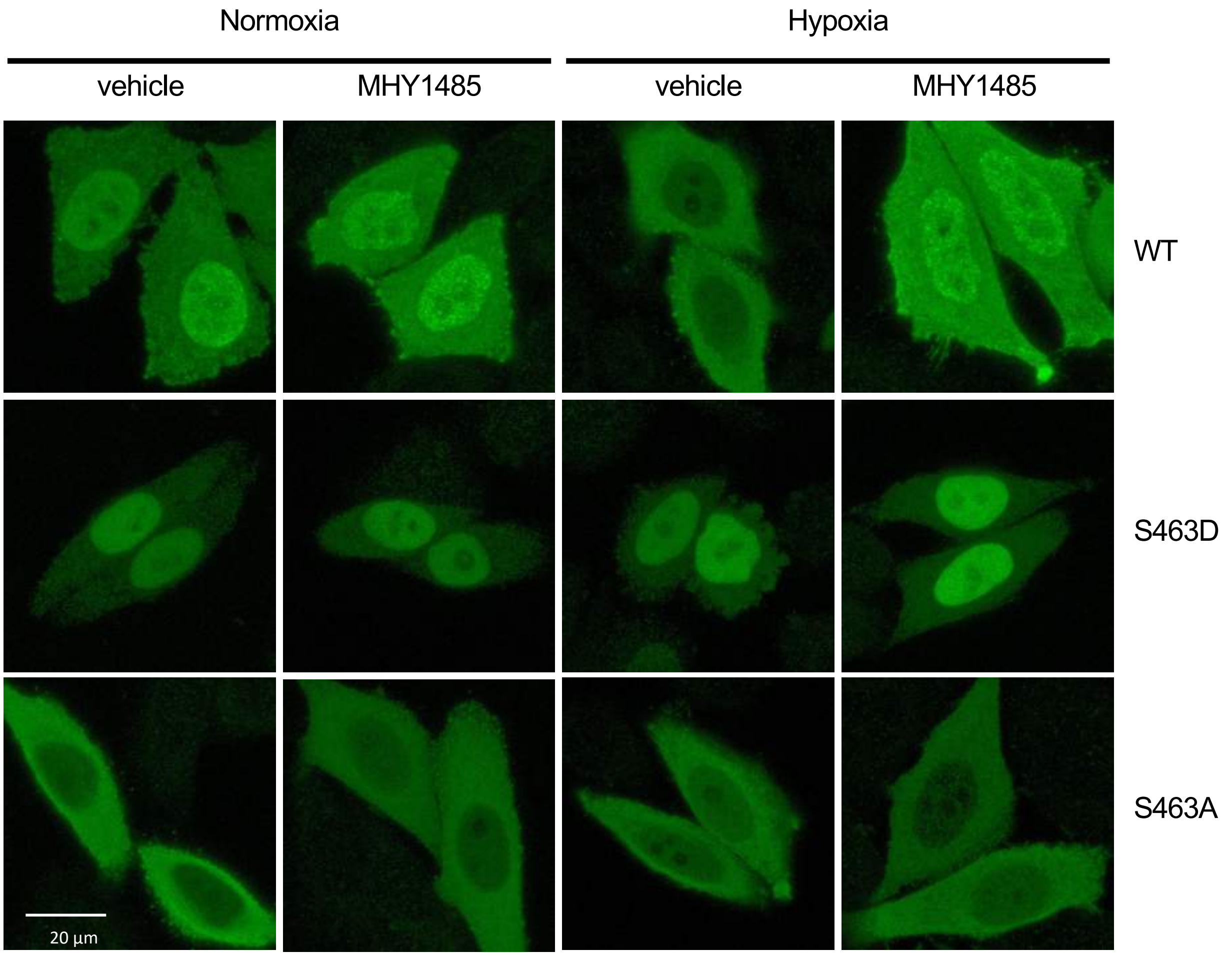
